# FaDA: A Shiny web application to accelerate common lab data analyses

**DOI:** 10.1101/2020.04.23.055673

**Authors:** Yodit Feseha, Quentin Moiteaux, Estelle Geffard, Gérard Ramstein, Sophie Brouard, Richard Danger

**Affiliations:** Université de Nantes, CHU Nantes, Inserm, Centre de Recherche en Transplantation et Immunologie, UMR 1064, ITUN, F-44000 Nantes, France; Labex IGO, Nantes, France; Université de Nantes, LS2N DUKe, UMR6004, Centrale Nantes, IMT Atlantique, INRIA and CNRS, Nantes, France; Centre d’Investigation Clinique en Biothérapie, Centre de ressources biologiques (CRB), Nantes, France

**Keywords:** Data Analysis, Programming, R, R Shiny, Gene Expression, Cytometry, Data Visualization, Computational Analysis

## Abstract

**Background:** Web-based data analysis and visualization tools are mostly designed for specific purposes, such as data from whole transcriptome RNA sequencing or single-cell RNA sequencing. However, limited efforts have been made to develop tools designed for data of common laboratory data for non-computational scientists. The importance of such web-based tool is stressed by the current increased samples capacity of conventional laboratory tools such as quantitative PCR, flow cytometry or ELISA.

**Results:** We provide a web-based application FaDA, developed with the R Shiny package providing users to perform statistical group comparisons, including parametric and non-parametric tests, with multiple testing corrections suitable for most of the standard wet-lab analyses. FaDA provides data visualization such as heatmap, principal component analysis (PCA) and receiver operating curve (ROC). Calculations are performed through the R language.

**Conclusions:** FaDA application provides a free and intuitive interface allowing biologists without bioinformatic skills to easily and quickly perform common lab data analyses. The application is freely accessible at https://shiny-bird.univ-nantes.fr/app/Fada

**Abbreviations:** AUC: Area Under the Curve; FaDA: Fast Data Analysis; GEO: Gene Expression Omnibus; ELISA: enzyme-linked immunosorbent assay; PCA: Principal Component Analysis; qPCR: quantitative PCR; ROC: Receiver Operating Curve.

## Background

Increasing web-based data analysis and visualization tools developed using the R programming Shiny package (1) are proposed to researchers. These tools are useful to analyze data from different perspectives and providing interactive visualizations. Hence, Shiny tools are enabling wet-laboratory researchers the ability to take advantage of bioinformatics advancements(2). While being free and saving time in the analytic stages without computational skills, most of the current online Shiny applications are dedicated to specific objectives or technologies such as *shinyheatmap* to generate heatmaps for large datasets(3), *shinyCircos* to build Circos plots from genomic data(4) or *shinyGEO* to analyze gene expression datasets directly from the Gene Expression Omnibus (GEO) repository(5). However, only a few of these applications have been designed for data generated from common laboratory technics such as quantitative PCR, flow cytometry or enzyme-linked immunosorbent assay (ELISA). The technological advances in these methods have allowed researchers to generate significant data output. Flow cytometry technologies run a high number of samples with a tenth of fluorochrome parameter combinations. Alongside, multiplex ELISAs produces a read of up to tenth cytokines per well and advancements in quantitative PCR (qPCR) devices allow analysis of samples in less than an hour. These high-data outputs leave laboratory researchers with time-consuming data analysis process. Furthermore, for such analyses, researchers usually perform targeted parameter analysis with several hands-on processes, increasing loss of information and human error risks.

We created a friendly-user and interactive web Shiny application supporting regular laboratory analyses from a wide array of data, including flow cytometry and qPCR data. This application allows researchers to perform classical statistical group comparisons, including parametric and non-parametric tests with multiple testing corrections and heatmap, principal component analysis (PCA), receiver operating curves (ROC) and correlogram visualizations. The FaDA application is freely accessible at https://shiny-bird.univ-nantes.fr/app/Fada

## Implementation

### FaDA Application

FaDA is developed in R programming language (release 3.6.1, http://www.rproject.org)(6) and implemented as a web application using the R Shiny package (version 1.4.0) from R Studio (http://shiny.rstudio.com). As an open-source application, the FaDA code is available through GitHub at https://github.com/danger-r/FaDAapp. FaDA is dockerized using the Docker software (https://www.docker.com/) and make available through ShinyProxy on a Linux server (CentOS 7 with 6 Go RAM allowed for FaDA) hosted at the genomic Bird core facility within the University of Nantes (https://pf-bird.univ-nantes.fr/). FaDA uses integrated work frames of R packages allowing an intuitive interface. A complete list of used packages may be found in supplementary table S1. The interface layout is built using the *shiny* and *shinythemes* packages with a sidebar for user interaction and with six main panels (*About, Tutorial, Data Analysis, Heatmap & PCA, Correlation* and *ROC curves*); with subtabs available within these panels (Figure 1).

**Figure 1:**
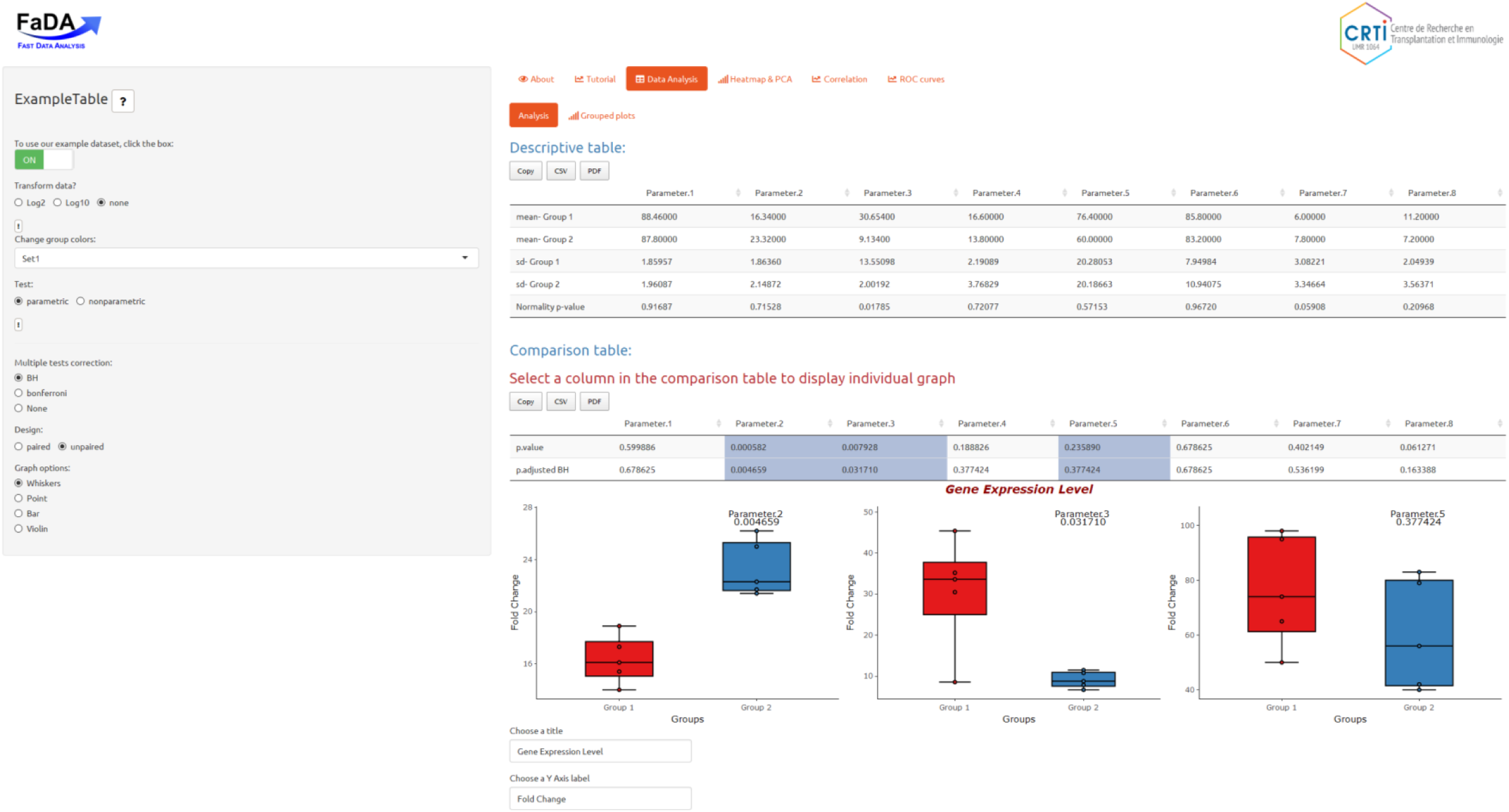
Overview Display of FaDA Application. Upon data uploading, users are automatically directed to the *Data Analysis* tabset (highlighted in orange) to view statistical summary of their dataset. The 6 different tabsets are available in the main panel (*About, Tutorial, Data analysis, Heatmap & PCA, Correlation*, and *ROC Curves*) while the sidebar display various options regarding data transformation (log10 or log2), statistical analysis (parametric or non-parametric, multiple correction options) and data visualisation options (group colors, graph options) depending on the selected tabset.

### Data Upload and File Input

The application starts with the *About* page, which displays general and background information of the application. The sidebar provides a simple demonstration dataset, including virtual data from 2 groups with 5 samples each, to explore the features of the web application. Alongside, data are up-loaded in a text format (tabular-delimited ‘.txt’ or ‘.csv’ file), with a point or a coma as a decimal separator. To allow flexible use of the application with a minimum preparation time; samples identification is in columns or rows. FaDA input only requires unique names of samples identification and the second row or column to be named “Group” to identify sample group labels. Users can find tutorial page displaying available tools FaDA provides.

### Statistical Summary

FaDA initially formulates a descriptive statistical summary with the upload of a dataset. The statistical analysis table presents mean and standard deviation values or median and interquartile interval (IQR) per group parameters; for parametric or nonparametric options, respectively. The p-value of the Shapiro-Wilk normality test indicates whether the distribution of the dataset differs from Gaussian distribution guiding the users toward parametric or nonparametric tests. The data can be transformed to log2 or log10, mainly useful for gene expression datasets. Group comparisons are performed using parametric t.test or ANOVA test with Tukey’s ‘Honest Significant Difference’ method for multiple groups comparisons. Alternatively, non-parametric group comparisons are available with the Mann Whitney test or the Kruskal Wallis test with Dunn’s test of multiple comparisons using the *FSA* package(7). In order to correct for false discovery rates due to multiple testing, statistical p-value corrections are performed with Bonferroni or Benjamini & Hochberg (BH) methods(8).

### Graph Visualization Plots

Shiny allows for built-in support of interactive graph plots of data using R’s graph representative and graph plot packages *gplots* and *ggplots2* (9). The graph plots available include box-and-whiskers, points, bars, and violins plots. Using the *plotly* package(10), interactive features are displayed, including zooming, panning, selecting, and downloading .png plots. Heatmap data representation is allowed either as static and interactive heatmaps. Static heatmap, may be customized using the *ComplexHeatmap* package(11), by adding sample hierarchical clustering and color schemes. PCA allows displaying covariance matrix and PCA plots to identify potential outliers or sample clustering. In the case of missing values, imputation is performed using the ten nearest neighbor averaging with the *impute* package(12). Both heatmap and PCA are visualized in an interactive mode, using the *heatmaply* and *plotly* packages, respectively(10,13).

### Correlation analysis

To assess the correlation between parameters, correlation coefficients are summarized in a correlogram thanks to the *corrplot* package(14). Individual correlation graphs display scatter plots of two selected parameters with the given correlation of these two parameters. Correlation coefficients (r) and statistical significance tests are calculated either with the parametric Pearson correlation or the Spearman non-parametric methods. Since complete observation are used to calculate correlation, the ten nearest neighbor averaging method is used to impute missing values(12).

### ROC Curves

Receiver Operating Characteristics (ROC) Curves and Area Under the Curve (AUC) can be viewed on the ROC curve tab using the *pROC* package(15).

## Results

### Case Study 1 – Gene expression data

We use a 20-gene expression dataset from peripheral blood from two groups of renal transplanted patients: 46 operational tolerant patients who stopped immunosuppressive regimen while keeping a stable renal function - and 266 renal transplanted patients with stable function under immunosuppression(16). This matrix was already normalized (mean-centred log-intensity values divided by standard deviation), so no transformation, *e.g.* log2 transformation, was applied. Given the gene expression matrix, FaDA allows clear discrimination of both population patients using heatmap and PCA visualization (Figure 2A-B). The first component of the PCA (PC1) explained 52 % of the observed variance. ROC curves analysis highlights individual genes able to discriminate both populations with AUCs above 0.7, such as the gene *AKR1C3* reaching an AUC of 0.796 (Figure 2C). The correlogram allows identifying correlated genes *MS4A1, CD22, CD79B, FCRL2, BLK* and *TCL1A* (figure 2D), in accordance with the previous signature found in operational tolerance and implication of B cells(17).

**Figure 2:**
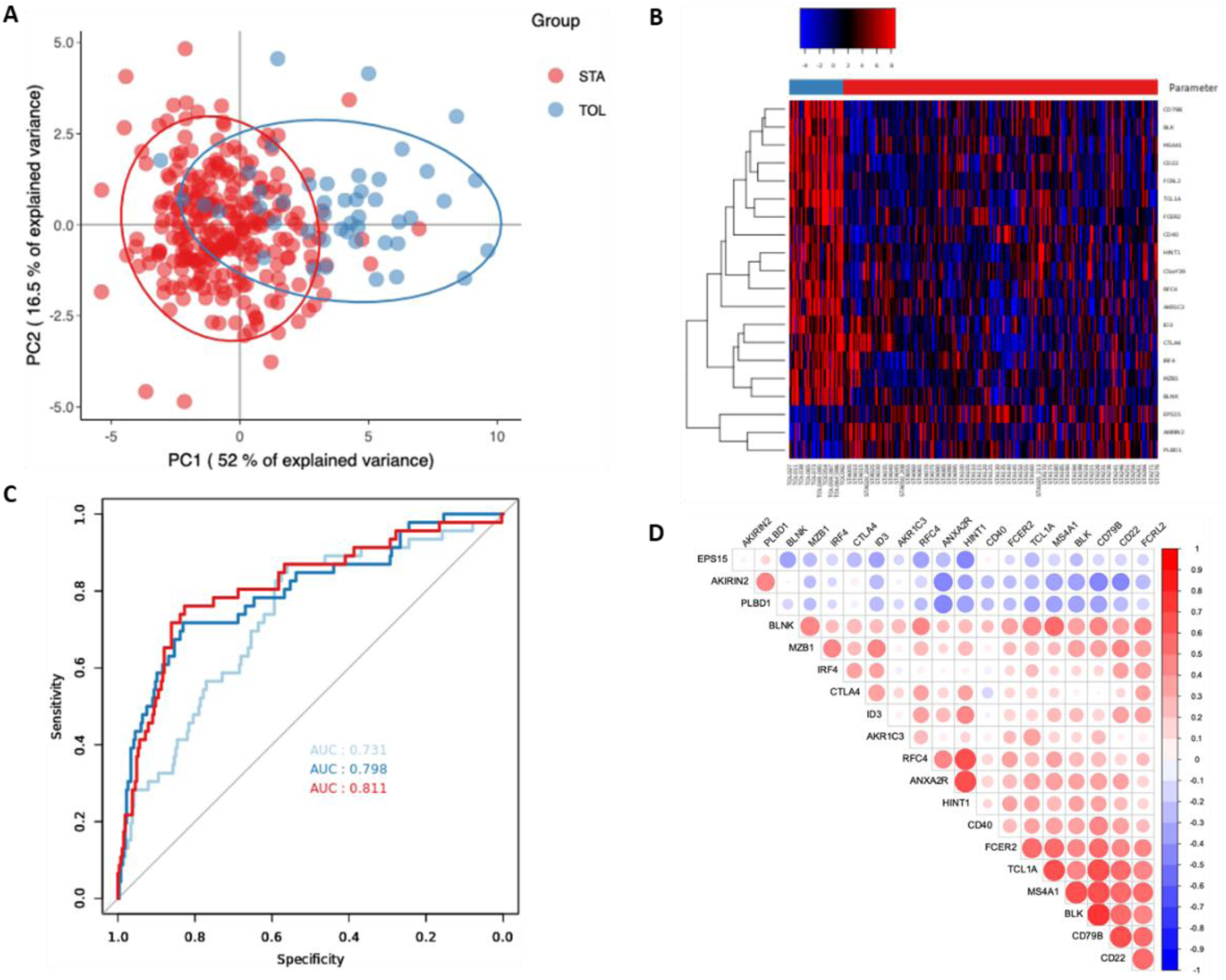
Data analysis of 20-gene expression dataset from renal transplanted patients. A) PCA and B) heatmap plots highlighting the clear gene expression difference of the two groups, TOL (blue) and STA (red). C) ROC curves of samples groups displaying AUC of the selected genes *AKR1C3, AKIRIN2* and *CD22.* D) Gene-gene correlation analysis using correlogram highlighting groups of genes.

### Case Study 2 – Flow Cytometry Data

We benefit from a previous study aimed to characterize circulating follicular T helper cells (cTfh) in peripheral blood of renal transplanted patients^15^. We reported on a notable impact of the anti-thymocyte globulin (ATG)-depleting induction treatment (n=87) compared to basiliximab non-depleting treatment (n=145) or the absence of induction therapy (n=5) on the frequency of total CD4^+^ lymphocytes and on activated cTfh subsets namely CXCR5^+^PD1^+^, CXCR5^+^PD1^+^ICOS^+^, CXCR5^+^PD1^+^CXCR3^-^ at one year after transplantation. Using FaDA, we can exhibit here, and as previously shown, that patients with depleting treatment exhibited lower levels of total CD4^+^ lymphocytes but higher frequencies of activated cTfh subsets using Benjamini-Hodchberg multi-testing correction (adjusted p.value <0.0001, figure 3A-B). Heatmap of dataset exhibits higher levels of activated cTfh subsets in depleting treatment groups (figure 3C).

**Figure 3:**
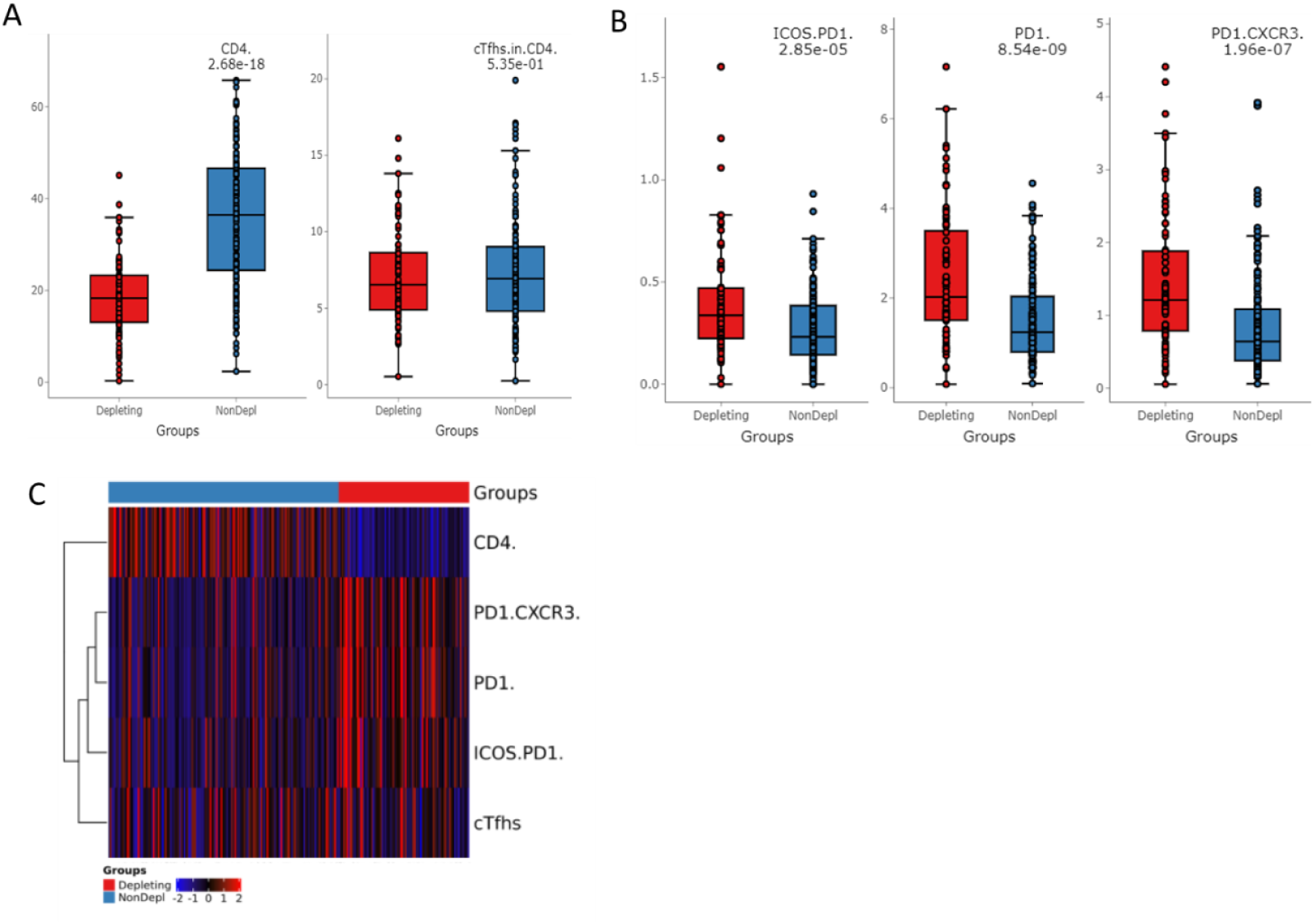
Cytometric dataset analysis using FaDA. A-B) Comparison at one-year post-transplantation of patients receiving ATG-depleting induction treatment (n=87) and basiliximab non-depleting treatment (n=145) or the absence of induction therapy (n=5) on the frequency of CD4^+^ cells, total cTfh^+^ and cTfh subsets, namely CXCR5^+^PD1^+^, CXCR5^+^PD1^+^ICOS^+^, CXCR5^+^PD1^+^CXCR3^-^. BH corrected p-values of t-tests are displayed. C) Heatmap graph represents a visual summary of flow cytometry data.

## Discussion

Here we demonstrate the easy and helpfulness use of FaDA application for the analysis of commonly generated data from flow cytometry and gene expression microarrays. FaDA web application is free and user-friendly, provided for non-computational scientists to easily and rapidly perform data analysis; while reducing the error arising from hands-on data analysis regularly used by wet-laboratory researchers. The FaDA application allows users to profit from various data visualization options given a glance of data analysis, identifying significant findings and possibly highlighting outliers with limited time consumption. We have used two cases study from previously published datasets(16,18) to exhibit the usefulness of FaDA for common data generated from non-computational researchers; that entails microarray and flow cytometry. Note that despite FaDA proposal of various statistical options, it cannot replace recommendations from a statistician that users may need for particular cases as for any analysis software. While we are offering user support, we planned to continue to develop this application providing additional tests and visualisation tools.

## Acknowledgments

We wish to thank the GenoBird Core Facility for their technical support and hosting the FaDA application.

## Authors’ Contributions

YF, QM and RD developed the application. YF and RD wrote the manuscript. EB, GR and SB reviewed the manuscript. All authors revised and approved the final manuscript.

## Supplementary Data

**Table S1:**
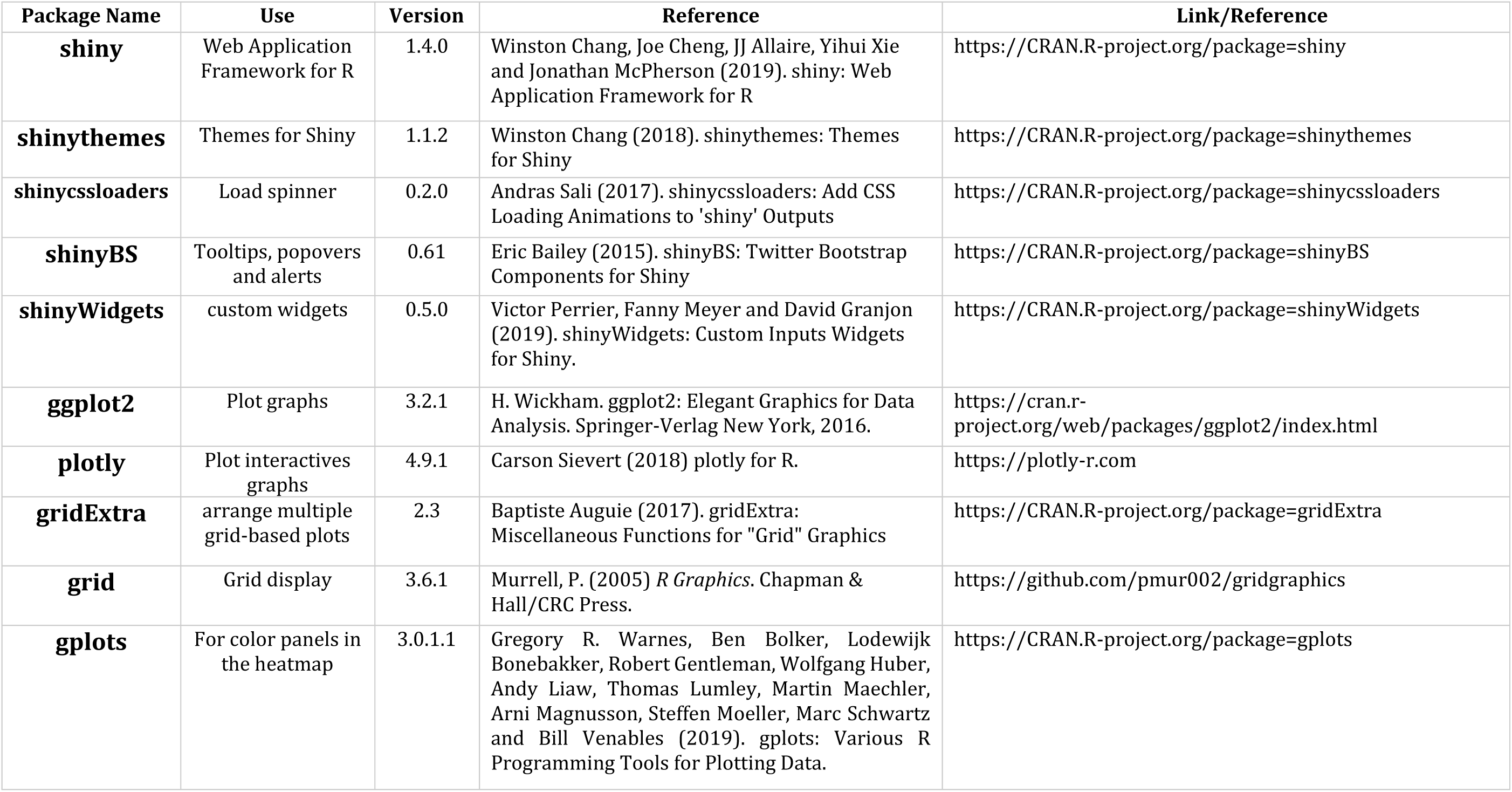
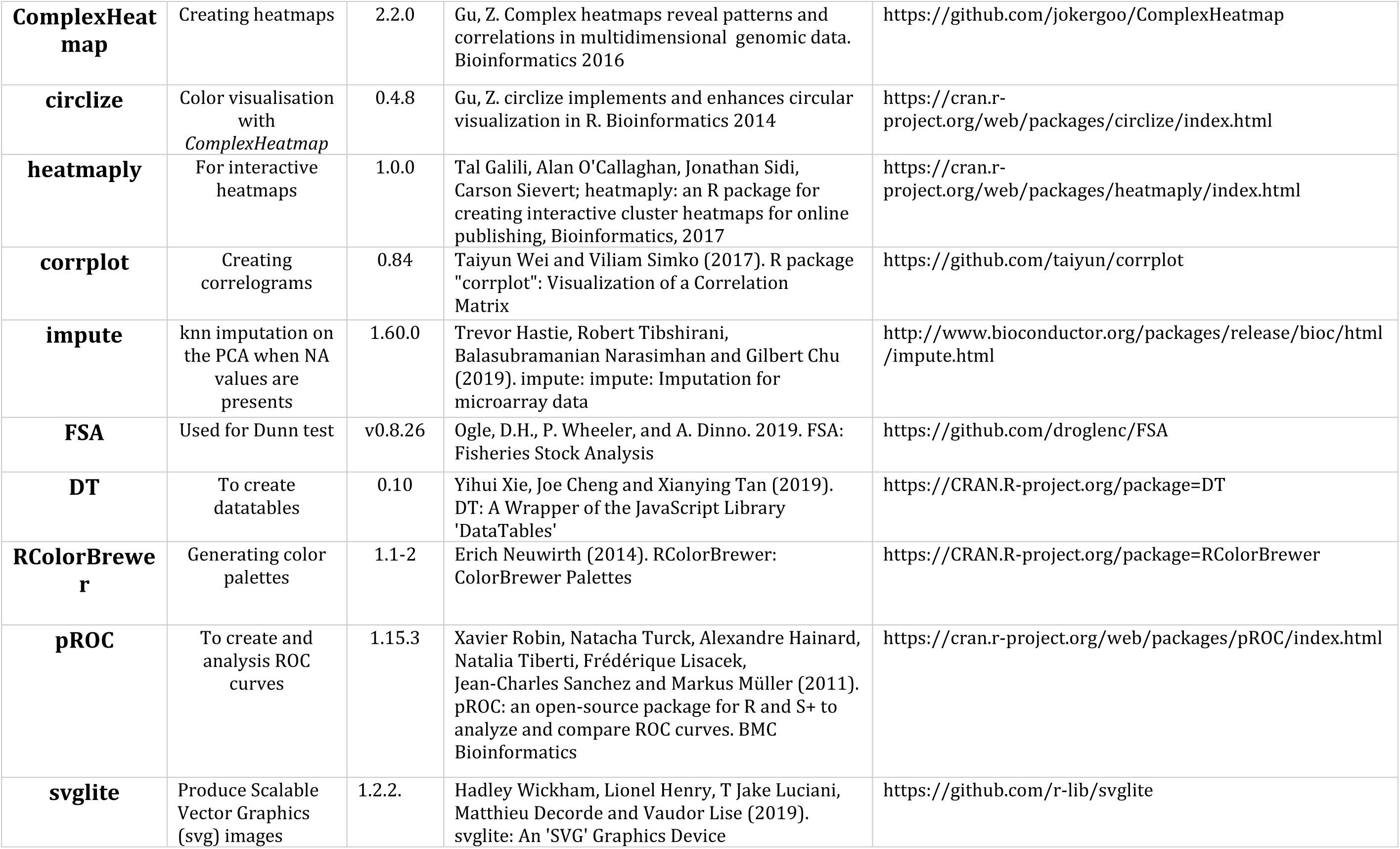
List of R/Shiny Packages

| Package Name | Use | Version | Reference | Link/Reference |
| --- | --- | --- | --- | --- |
| **shiny** | Web Application Framework for R | 1.4.0 | Winston Chang, Joe Cheng, JJ Allaire, Yihui Xie and Jonathan McPherson (2019). shiny: Web Application Framework for R | https://CRAN.R-project.org/package=shiny |
| **shinythemes** | Themes for Shiny | 1.1.2 | Winston Chang (2018). shinythemes: Themes for Shiny | https://CRAN.R-project.org/package=shinythemes |
| **shinycssloaders** | Load spinner | 0.2.0 | Andras Sali (2017). shinycssloaders: Add CSS Loading Animations to 'shiny' Outputs | https://CRAN.R-project.org/package=shinycssloaders |
| **shinyBS** | Tooltips, popovers and alerts | 0.61 | Eric Bailey (2015). shinyBS: Twitter Bootstrap Components for Shiny | https://CRAN.R-project.org/package=shinyBS |
| **shinyWidgets** | custom widgets | 0.5.0 | Victor Perrier, Fanny Meyer and David Granjon (2019). shinyWidgets: Custom Inputs Widgets for Shiny. | https://CRAN.R-project.org/package=shinyWidgets |
| **ggplot2** | Plot graphs | 3.2.1 | H. Wickham. ggplot2: Elegant Graphics for Data Analysis. Springer-Verlag New York, 2016. | https://cran.r-project.org/web/packages/ggplot2/index.html |
| **plotly** | Plot interactives graphs | 4.9.1 | Carson Sievert (2018) plotly for R. | https://plotly-r.com |
| **gridExtra** | arrange multiple grid-based plots | 2.3 | Baptiste Auguie (2017). gridExtra: Miscellaneous Functions for "Grid" Graphics | https://CRAN.R-project.org/package=gridExtra |
| **grid** | Grid display | 3.6.1 | Murrell, P. (2005) *R Graphics*. Chapman & Hall/CRC Press. | https://github.com/pmur002/gridgraphics |
| **gplots** | For color panels in the heatmap | 3.0.1.1 | Gregory R. Warnes, Ben Bolker, Lodewijk Bonebakker, Robert Gentleman, Wolfgang Huber, Andy Liaw, Thomas Lumley, Martin Maechler, Arni Magnusson, Steffen Moeller, Marc Schwartz and Bill Venables (2019). gplots: Various R Programming Tools for Plotting Data. | https://CRAN.R-project.org/package=gplots |
| **ComplexHeatmap** | Creating heatmaps | 2.2.0 | Gu, Z. Complex heatmaps reveal patterns and correlations in multidimensional genomic data. Bioinformatics 2016 | https://github.com/jokergoo/ComplexHeatmap |
| **circlize** | Color visualisation with *ComplexHeatmap* | 0.4.8 | Gu, Z. circlize implements and enhances circular visualization in R. Bioinformatics 2014 | https://cran.r-project.org/web/packages/circlize/index.html |
| **heatmaply** | For interactive heatmaps | 1.0.0 | Tal Galili, Alan O'Callaghan, Jonathan Sidi, Carson Sievert; heatmaply: an R package for creating interactive cluster heatmaps for online publishing, Bioinformatics, 2017 | https://cran.r-project.org/web/packages/heatmaply/index.html |
| **corrplot** | Creating correlograms | 0.84 | Taiyun Wei and Viliam Simko (2017). R package "corrplot": Visualization of a Correlation Matrix | https://github.com/taiyun/corrplot |
| **impute** | knn imputation on the PCA when NA values are presents | 1.60.0 | Trevor Hastie, Robert Tibshirani, Balasubramanian Narasimhan and Gilbert Chu (2019). impute: impute: Imputation for microarray data | http://www.bioconductor.org/packages/release/bioc/html/impute.html |
| **FSA** | Used for Dunn test | v0.8.26 | Ogle, D.H., P. Wheeler, and A. Dinno. 2019. FSA: Fisheries Stock Analysis | https://github.com/droglenc/FSA |
| **DT** | To create datatables | 0.10 | Yihui Xie, Joe Cheng and Xianying Tan (2019). DT: A Wrapper of the JavaScript Library 'DataTables' | https://CRAN.R-project.org/package=DT |
| **RColorBrewer** | Generating color palettes | 1.1-2 | Erich Neuwirth (2014). RColorBrewer: ColorBrewer Palettes | https://CRAN.R-project.org/package=RColorBrewer |
| **pROC** | To create and analysis ROC curves | 1.15.3 | Xavier Robin, Natacha Turck, Alexandre Hainard, Natalia Tiberti, Frédérique Lisacek, Jean-Charles Sanchez and Markus Müller (2011). pROC: an open-source package for R and S+ to analyze and compare ROC curves. BMC Bioinformatics | https://cran.r-project.org/web/packages/pROC/index.html |
| **svglite** | Produce Scalable Vector Graphics (svg) images | 1.2.2. | Hadley Wickham, Lionel Henry, T Jake Luciani, Matthieu Decorde and Vaudor Lise (2019). svglite: An 'SVG' Graphics Device | https://github.com/r-lib/svglite |

